# An exon junction complex-independent function of Barentsz in neuromuscular synapse growth

**DOI:** 10.1101/2021.02.13.430688

**Authors:** Cheuk Hei Ho, Jean-Yves Roignant, Zuojian Tang, Stuart Brown, Jessica E. Treisman

**Affiliations:** Skirball Institute for Biomolecular Medicine and Department of Cell Biology; Center for Health Informatics and Bioinformatics, NYU Langone Medical Center, 540 First Avenue, New York, NY 10016; Center for Integrative Genomics, Génopode Building, Faculty of Biology and Medicine, University of Lausanne, CH-1015, Lausanne, Switzerland; Computational Biology at Ridgefield US, Global Computational Biology and Digital Science, Boehringer Ingelheim; ExxonMobil Corporate Strategic Research, Annandale, NJ

**Keywords:** exon junction complex, Barentsz, neuromuscular junction, Dawdle, synapse

## Abstract

The exon junction complex controls the translation, degradation and localization of spliced mRNAs, and three of its four core subunits also play a role in splicing. Here we show that the fourth subunit, Barentsz, has distinct biological functions within and separate from the exon junction complex in neuromuscular development. Barentsz controls the distribution of mitochondria in larval muscles, a function that also depends on other subunits of the exon junction complex and that is not rescued by a transgene in which residues required for binding to the core subunit eIF4AIII are mutated. In contrast, interactions with the exon junction complex are not required for Barentsz to promote the growth of neuromuscular synapses. We found that the Activin ligand Dawdle shows reduced expression in *barentsz* mutants and acts downstream of Barentsz to control synapse growth. Both *barentsz* and *dawdle* are required in motor neurons, muscles and glia for normal synapse growth, and exogenous Dawdle can rescue synapse growth in the absence of *barentsz*. These results identify a biological function for Barentsz that is independent of the exon junction complex.

## Introduction

Post-transcriptional regulation enables rapid and localized changes in gene expression, making it an important regulatory mechanism in the nervous system (Goldie and Cairns, 2012). The exon junction complex (EJC) links distinct modes of post-transcriptional regulation by marking properly spliced mRNAs for preferential translation, efficient nonsense-mediated decay (NMD), or specific subcellular localization (Boehm et al., 2014; Chang et al., 2007; Chazal et al., 2013; Choe et al., 2014; Ghosh et al., 2012; Ma et al., 2008; Metze et al., 2013; Nott et al., 2004; Palacios et al., 2004; Schlautmann and Gehring, 2020). During splicing, the core EJC protein eIF4AIII (EIF4A3 in humans) is recruited by the spliceosomal protein CWC22 to mRNA 20-24 nucleotides upstream of exon junctions (Alexandrov et al., 2012; Barbosa et al., 2012; Gehring et al., 2009; Steckelberg et al., 2012). A dimer of Mago nashi (Mago)/MAGOH and Y14/RBM8A then associates with eIF4AIII (Gehring et al., 2009; Herold et al., 2009). These proteins together with the accessory subunit RnpS1/RNPS1 facilitate the splicing of a subset of introns that includes introns in *MAP kinase, piwi*, and pro-apoptotic transcripts (Ashton-Beaucage et al., 2010; Haremaki and Weinstein, 2012; Hayashi et al., 2014; Malone et al., 2014; Michelle et al., 2012; Roignant and Treisman, 2010). In addition, this complex prevents the splicing of cryptic splice sites within introns (Blazquez et al., 2018; Boehm et al., 2018; Gehring and Roignant, 2020).

A fourth core subunit, known as Barentsz (Btz), Cancer susceptibility candidate gene 3 (CASC3), or Metastatic lymph node 51 (MLN51), associates with the complex following the completion of splicing, and is required for the effects of the EJC on translation, NMD and mRNA localization (Chazal et al., 2013; Palacios et al., 2004; Shibuya et al., 2006; van Eeden et al., 2001). The transition between RNPS1 and CASC3-containing forms of the EJC alters the set of transcripts subject to NMD, although distinct effects have been reported in two studies (Gerbracht et al., 2020; Mabin et al., 2018). In the mouse brain, haploinsufficiency for *Magoh, Rbm8a* or *Eif4a3* causes severe microcephaly, but complete loss of *Casc3* has a much milder effect that can be attributed to developmental delay (Mao et al., 2017; Mao et al., 2016; Mao et al., 2015; Silver et al., 2010). Btz/CASC3 shuttles between the cytoplasm and the nucleus, and can localize to stress granules, P-bodies and neuronal RNP granules independently of other EJC subunits (Baguet et al., 2007; Barbee et al., 2006; Cougot et al., 2014; Fritzsche et al., 2013; Macchi et al., 2003; Vessey et al., 2006). However, a biological function for Btz/CASC3 acting outside the EJC has not yet been reported.

During development, the synapses formed by motor neurons on their target muscles, known as neuromuscular junctions (NMJs), add new branches and synaptic boutons as the muscles increase in size (Menon et al., 2013; Schuster et al., 1996). This coordinated growth requires multiple signals to pass between the two cell types. Motor neurons secrete Wingless (Wg), which acts both on the nerve terminals through its canonical signaling pathway, and on the muscle through a novel mechanism involving nuclear import of the Frizzled receptor (Koles and Budnik, 2012; Mathew et al., 2005; Miech et al., 2008; Mosca and Schwarz, 2010). Muscles secrete the retrograde signal Glass bottom boat (Gbb), a Bone morphogenetic protein (BMP) family member that acts on motor neurons through the Wishful thinking (Wit) receptor, leading to expression of the exchange factor Trio and enhanced function of the receptor protein tyrosine phosphatase Lar (Ball et al., 2010; Berke et al., 2013; Keshishian and Kim, 2004). Gbb is also secreted from motor neurons in an activity-dependent manner in association with the ectodomain of the Crimpy protein, and this complex has a distinct function in regulating neurotransmitter release (James et al., 2014). Gbb production by muscles is dependent on two Activin-like ligands, Dawdle (Daw) and Maverick (Mav), which are produced by glial cells surrounding the nerve terminal as well as by the muscles themselves (Ellis et al., 2010; Fuentes-Medel et al., 2012; Parker et al., 2006; Serpe and O’Connor, 2006). However, the canonical Activin signaling pathway does not appear to be necessary for NMJ growth (Kim and O’Connor, 2014). Additional signals also contribute to the control of synapse growth, including neurotrophins, stress response mechanisms, and a pathway that transmits information about muscle size through the extent of postsynaptic differentiation (Chen and Ganetzky, 2012; Ho and Treisman, 2020; Milton et al., 2011; Xiong et al., 2010).

Here we show that Btz has both EJC-dependent and EJC-independent functions in neuromuscular development. It acts in muscles to control the distribution of mitochondria, and this function requires physical and functional interactions with other EJC subunits. Btz also acts in motor neurons, muscles and glia to promote NMJ growth, and interaction with other EJC subunits is dispensable for this process. Analysis of transcripts that show altered levels in the central nervous system or larval carcass in *btz* mutants identified changes in expression of the Activin family member *daw*, as well as of two potential regulators of Activin signaling. *daw* acts in the same tissues as *btz* to promote NMJ growth, and exogenous *daw* can rescue NMJ size in *btz* mutants. These results indicate that Btz acts independently of the EJC to stimulate neuromuscular synapse growth, and that this function is mediated by Activin signaling.

## Results

### A tool for the analysis of EJC-dependent and independent functions of Btz

We and others have previously demonstrated that the Mago, Y14 and eIF4AIII subunits of the EJC have a function in splicing that is independent of Btz (Ashton-Beaucage et al., 2010; Blazquez et al., 2018; Boehm et al., 2018; Haremaki and Weinstein, 2012; Hayashi et al., 2014; Malone et al., 2014; Michelle et al., 2012; Roignant and Treisman, 2010). To test whether Btz can also act independently of the other EJC subunits, we generated two rescue transgenes that express a GFP-tagged form of Btz with 5’ and 3’ endogenous regulatory sequences encompassing the entire intergenic region between *btz* and the adjacent genes (Fig. 1A). One transgene expresses the wild-type Btz protein (*Btz-WT*), while the other expresses a Btz protein with mutations in two conserved residues (H215A and D216A) in the SELOR (Speckle localizer and RNA-binding) module (*Btz-HD*). These changes correspond to the H220A and D221A mutations that have been shown to disrupt the association of human MLN51 with eIF4AIII and the EJC without preventing it from interacting with RNA (Ballut et al., 2005; Bono et al., 2006; Daguenet et al., 2012; Gehring et al., 2009). Both proteins were equally stable when expressed in cultured S2 cells (Fig. 1B).

**Figure 1:**
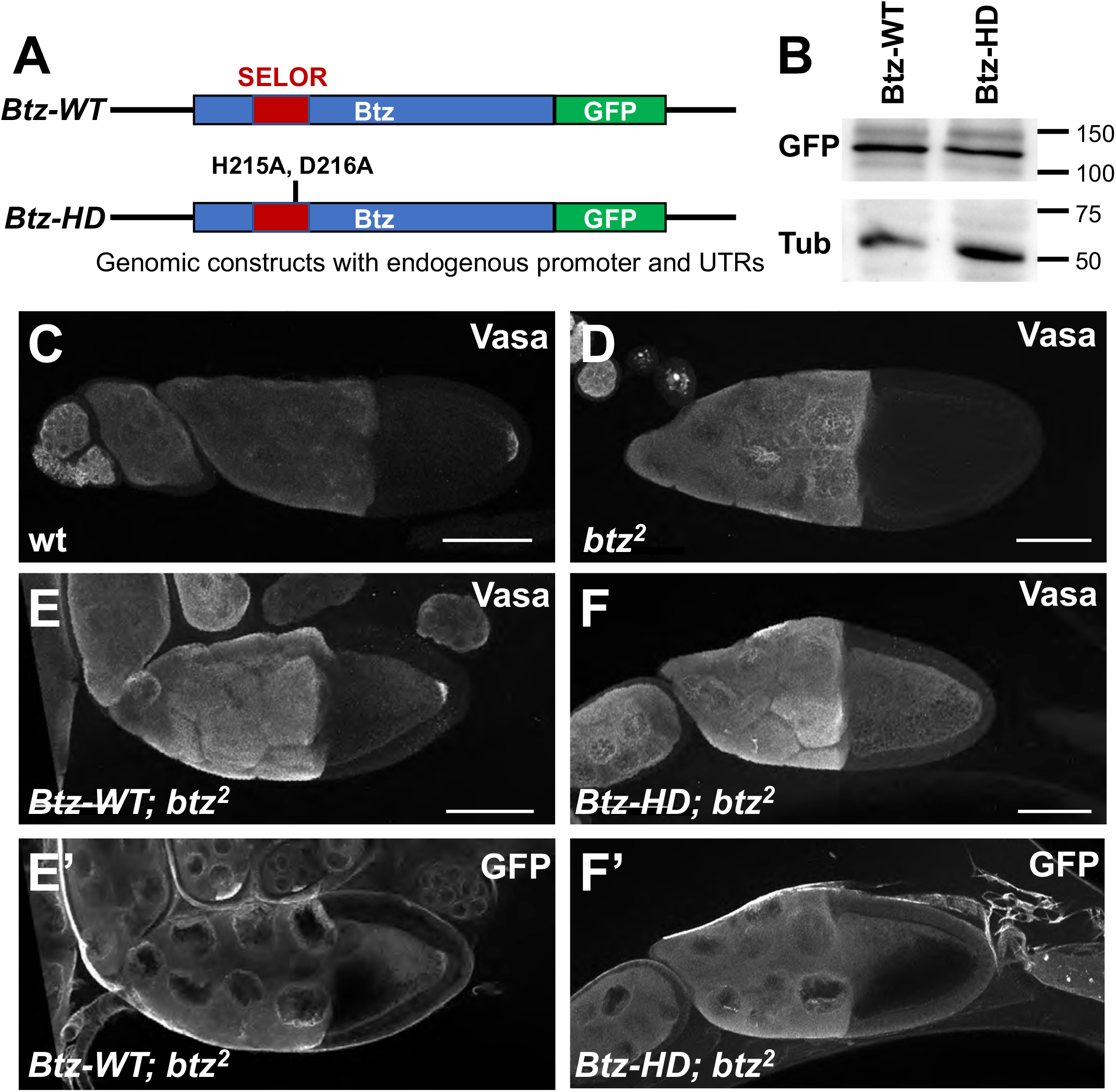
Interaction with the EJC is required for Btz function in the ovary. (A) diagram of the genomic rescue constructs in which wild type Btz (*Btz-WT*) or Btz with the H215A and D216A mutations (*Btz-HD*) is C-terminally tagged with GFP and driven by its endogenous regulatory sequences. (B) Western blot of S2 cell extracts transfected with the *Btz-WT* or *Btz-HD* plasmids, blotted with anti-GFP and anti-β-Tubulin (Tub). The H215A/D216A mutations do not destabilize the Btz protein. (C-F) show stage 10 egg chambers stained with anti-Vasa (C-F) or anti-GFP (E’, F’). (C) wild type; (D) *btz*^*2*^ germline clones; (E) *Btz-WT; btz*^*2*^ germline clones; (F) *Btz-HD; btz*^*2*^ germline clones. Posterior is to the right. Scale bars, 80 µm. Vasa localization to the posterior pole of the oocyte is lost in *btz* mutants and is rescued by the wild type but not the EJC interaction-defective transgene. Wild-type Btz-GFP, but not the mutant form, is also slightly enriched at the posterior pole.

The full EJC, including Btz, is required to localize *oskar* mRNA to the posterior of the oocyte, leading to the assembly of germ plasm there (Palacios et al., 2004; van Eeden et al., 2001). Therefore, we tested the ability of the wild-type and mutant transgenes to rescue posterior germ plasm formation in *btz* mutants, using Vasa protein as a readout for germ plasm (Breitwieser et al., 1996). As predicted, the the *Btz-WT* transgene rescued the loss of posteriorly localized Vasa in *btz* mutant oocytes, and the encoded Btz-WT-GFP protein was enriched at the posterior pole (Fig. 1C-E). However, Btz-HD-GFP failed to localize to the posterior pole or to rescue Vasa localization (Fig. 1F), confirming that H215 and D216 are necessary for Btz to function in the context of the EJC. These transgenes thus enable us to distinguish between EJC-dependent and possible EJC-independent functions of Btz.

### Btz acts through the EJC to control the distribution of mitochondria in muscle

In addition to its effect on *oskar* mRNA localization in the developing oocyte (Palacios et al., 2004; van Eeden et al., 2001), *btz* also has an essential role during development: most flies transheterozygous for a null allele of *btz* (van Eeden et al., 2001) and a deficiency covering the region (*Df(3R)BSC497*) die as late pupae. Our wild-type transgene fully rescued the lethality of these transheterozygous *btz* mutants, while the mutant transgene provided a partial rescue (Table 1), suggesting that some of the functions of Btz that are required for viability may be EJC-independent. We found that expressing a GFP-tagged form of Btz with a muscle-specific GAL4 driver fully rescued the mutant flies to viability, while expressing it with a motor neuron driver or a glial driver partially rescued the lethality (Table 1). These results indicate that the most critical requirement for Btz is in muscle, but it also functions in motor neurons and glia.

**Table 1:** Rescue of viability in *btz* mutants. The table shows the percentage of pupae from which adult flies eclosed and the total number of pupae counted for the indicated genotypes. Adult viability of *btz*^*2*^*/Df(3R)BSC497* is substantially rescued by the wild-type *btz* transgene and by *UAS-btz* expression ubiquitously or in muscles, and partially rescued by the mutant transgene and by *UAS-btz* expression in motor neurons or glia or by ubiquitous or muscle-specific *UAS-daw* expression.

| Genotype | % of eclosed pupae | n |
| --- | --- | --- |
| *btz/+* | 100 | 186 |
| *btz/Df* | 8.5 | 176 |
| *btz/Df, Btz<sup>WT</sup>* | 94.0 | 133 |
| *btz/Df, Btz<sup>HD</sup>* | 40.9 | 115 |
| *btz/Df, UAS-BTZ-GFP* | 6.5 | 92 |
| *btz/Df, BG380-GAL4>* | 7.4 | 95 |
| *btz/Df, BG380-GAL4>UAS-BTZ-GFP* | 24.3 | 103 |
| *btz/Df, repo-GAL4>* | 1.0 | 100 |
| *btz/Df, repo-GAL4>UAS-BTZ-GFP* | 24.1 | 83 |
| *btz/Df, mhc-Gal4>* | 6.9 | 131 |
| *btz/Df, mhc-Gal4>UAS-BTZ-GFP* | 88.6 | 220 |
| *btz/Df, mhc-Gal4>UAS-dawdle* | 28.5 | 165 |
| *btz/Df, da-GAL4>* | 5.4 | 74 |
| *btz/Df, da-GAL4>UAS-BTZ-GFP* | 97.4 | 78 |
| *btz/Df, da-GAL4>UAS-dawdle* | 23.5 | 132 |

We observed that one striking effect of loss of *btz* in larval muscles is a change in the distribution of mitochondria. In wild-type muscles, mitochondria detected either by staining for the ATP5A subunit of the ATP synthase complex or by electron microscopy are enriched at the dorsal surface, but also distributed between the muscle fibers (Fig. 2A, B, L, M). In *btz* mutants, almost all mitochondria are concentrated at the dorsal surface (Fig. 2C, N). We quantified these differences by measuring the ratio of anti-ATP5A staining intensity in the dorsal peak relative to the next largest peak in a line scan through the muscle (Fig. 2O). The normal distribution of ATP5A staining was restored by expressing *UAS-BtzGFP* in muscle with *G14-GAL4* or by the presence of our *Btz-WT* transgene (Fig. 2D, E, O), but not by the *Btz-HD* transgene (Fig. 2F, O), suggesting that this function of Btz is mediated by the EJC. To confirm this, we used the muscle-specific driver *C57-GAL4* to express UAS-RNAi transgenes targeting Btz or the EJC subunits Mago and eIF4AIII, and examined the localization of ATP5A staining. Depletion of each of these subunits caused mitochondria to concentrate at the dorsal surface (Fig. 2G-J, O). The specificity of the ATP5A antibody was confirmed by the lack of staining in muscles in which *bellwether* (*blw*), which encodes ATP5A, was knocked down (Fig. 2K). These results indicate that Btz acts as a subunit of the EJC to promote the even distribution of mitochondria within muscles.

**Figure 2:**
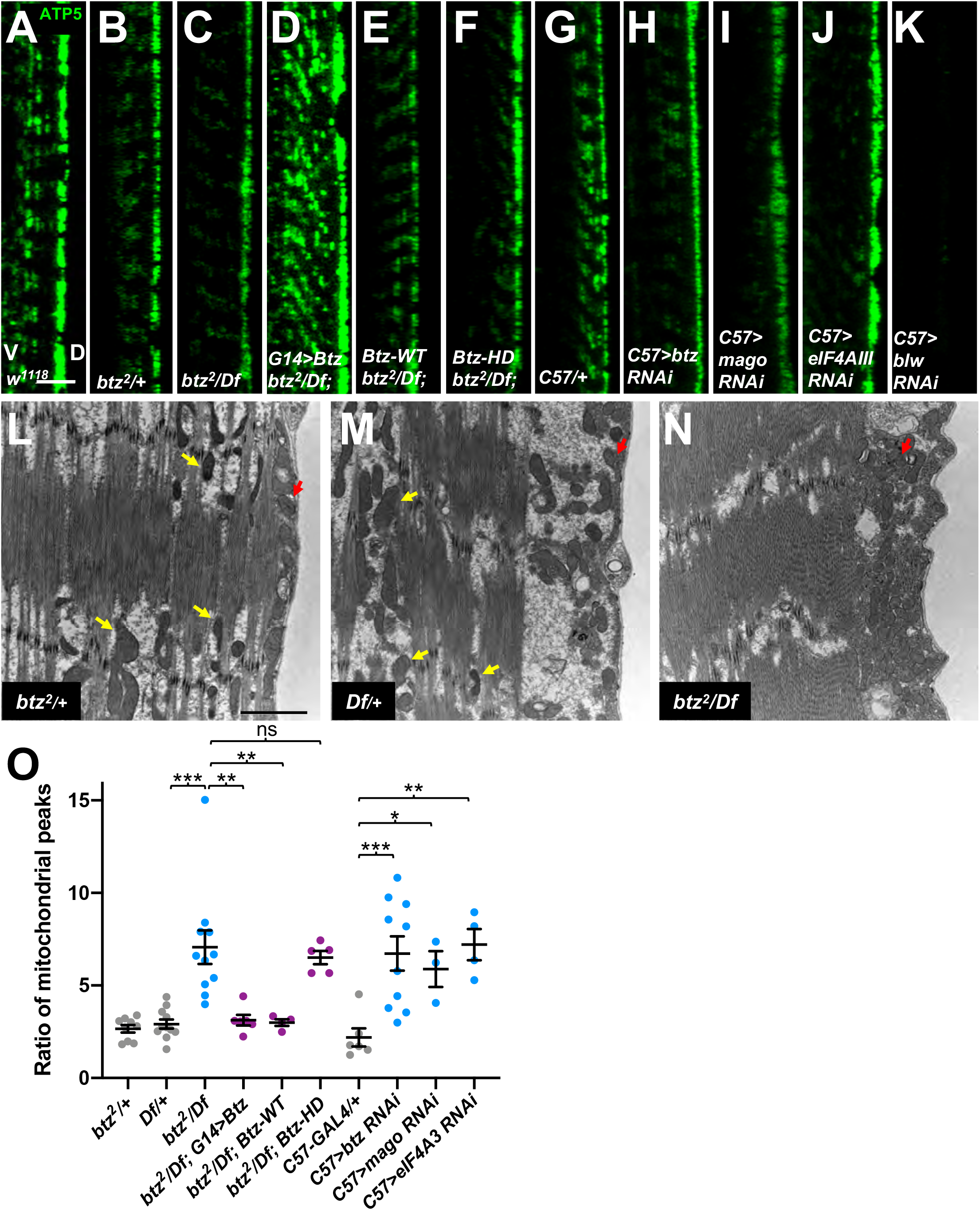
Btz acts as an EJC component to control the distribution of mitochondria in muscle. (A-K) Confocal xz-sections of larval muscle 6 in segment A3 stained with anti-ATP5A to label mitochondria. The dorsal surface (D) is to the right and ventral (V) to the left. (A) *w*^*1118*^; (B) *btz*^*2*^*/+*; (C) *btz*^*2*^*/ Df(3R)BSC497*; (D) *G14-GAL4/UAS-btz; btz*^*2*^*/ Df(3R)BSC497*; (E) *Btz-WT; btz*^*2*^*/ Df(3R)BSC497*; (F) *Btz-HD; btz*^*2*^*/ Df(3R)BSC497*; (G) *C57-GAL4/+*; (H) *C57-GAL4; UAS-btz RNAi*; (I) *C57-GAL4; UAS-mago RNAi*; (J) *C57-GAL4; UAS-eIF4AIII RNAi*; (K) *C57-GAL4; UAS-blw RNAi*. In wild type muscles, mitochondria are present at the dorsal surface and also distributed between the fibers, while in *btz* mutant muscles and in muscles expressing *btz, mago* or *eIF4AIII* RNAi they are concentrated at the dorsal surface. Expression of *UAS-btz* in muscle or the presence of the wild type, but not the HD mutant, *btz* transgene restores the wild type distribution to *btz* mutants. The loss of ATP5A staining in muscles in which *blw* is knocked down confirms that the signal is specific. (L-N) electron micrographs showing cross-sections of larval muscle 6 in segment A3, with the dorsal surface to the right. (L) *btz*^*2*^*/+*; (M) *Df(3R)BSC497/+*; (N) *btz*^*2*^*/ Df(3R)BSC497*. Yellow arrows indicate examples of mitochondria interspersed between muscle fibers, and red arrows indicate mitochondria at the dorsal surface. Scale bars, 10 µm (A-K) or 2 µm (L-N). (O) quantification of the ratio of the dorsal surface peak of ATP5 intensity to the second highest peak for the genotypes indicated. ***, p<0.001; **p<0.01; *p<0.05; ns, not significant by unpaired t-test. n=9 (*btz/+*), n=11 (*Df/+, btz/Df*), n=6 (*btz/Df; G14>btz, C57-GAL4/+*), n=4 (*btz/Df; Btz-WT, C57>eIF4AIII RNAi*), n=5 (*btz/Df; Btz-HD*), n=10 (*C57>btz RNAi*), or n=3 (*C57>mago RNAi*).

### Btz acts in motor neurons, muscles and glia to promote normal NMJ growth

Because *btz* acts in motor neurons and glia as well as muscles to promote viability, we considered the possibility that it might be required for normal development of the NMJ, a structure that is formed by the conjunction of these three cell types and influenced by signaling interactions between them. Indeed, we found that *btz* mutant NMJs had fewer synaptic boutons relative to their muscle surface area than controls (Fig. 3A-C, K). We confirmed that this reduction in bouton number was due to loss of *btz* function, because it could be rescued by ubiquitous expression of our *UAS-Btz-GFP* transgene with the *daughterless* (*da)-GAL4* driver (Fig. 3D, K).

**Figure 3:**
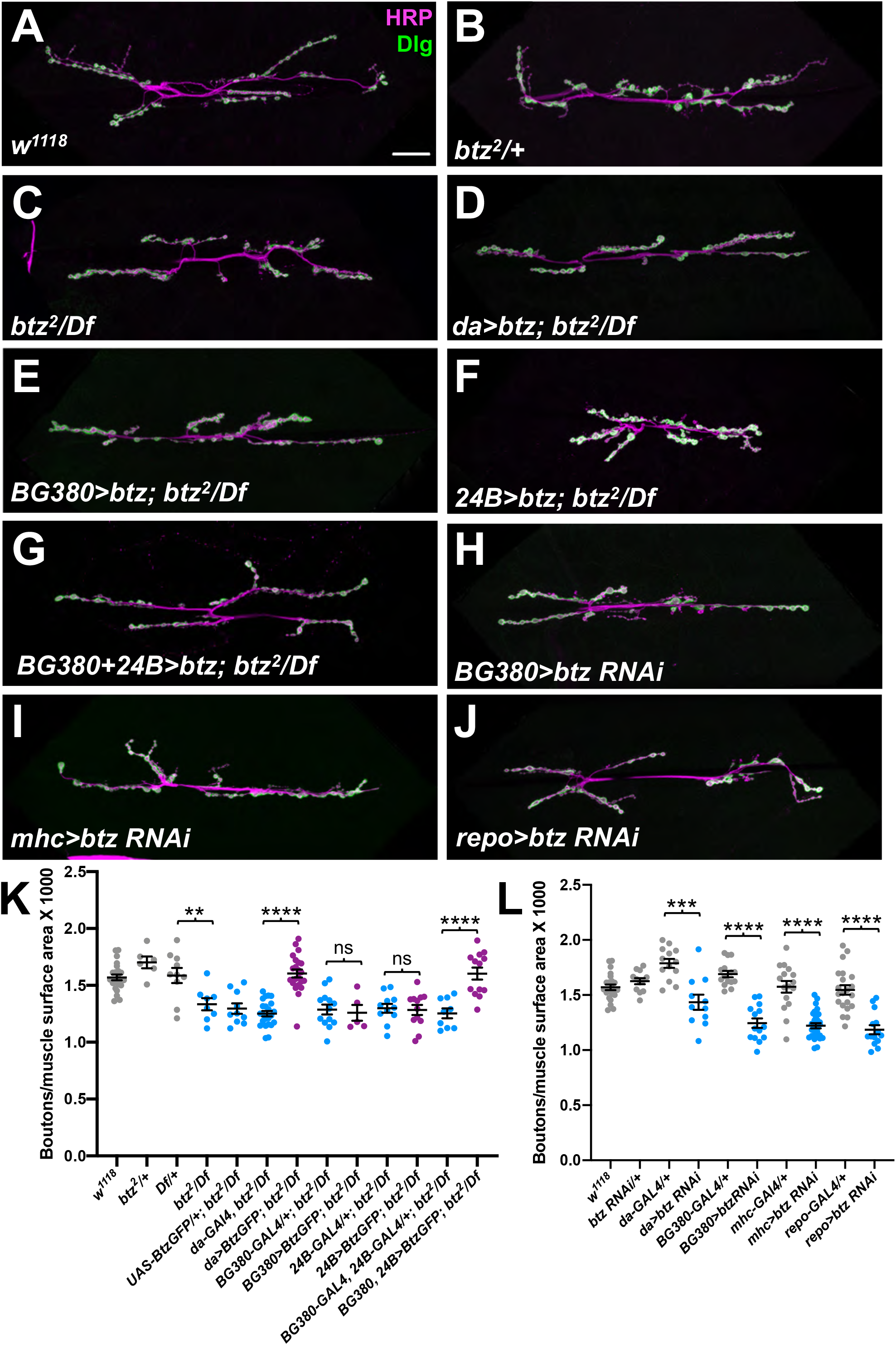
Btz acts in muscles, motor neurons and glia to control NMJ size. (A-J) confocal images of the NMJ on larval muscles 6 and 7 in segment A3, stained with anti-HRP (magenta) to label the nerve and anti-Dlg (green) as a postsynaptic marker of synaptic boutons. (A) *w*^*1118*^; (B) *btz*^*2*^*/+*; (C) *btz*^*2*^*/Df(3R)BSC497*; (D) *UAS-btz/+; da-GAL4, btz*^*2*^*/Df(3R)BSC497*; (E) *BG380-GAL4/+; UAS-btz/+; btz*^*2*^*/Df(3R)BSC497*; (F) *24B-GAL4/UAS-btz; btz*^*2*^*/Df(3R)BSC497*; (G) *BG380-GAL4/+; 24B-GAL4/UAS-btz; btz*^*2*^*/Df(3R)BSC497*; (H) *BG380-GAL4/+; UAS-btz RNAi/+*; (I) *mhc-GAL4/UAS-btz RNAi*; (J) *UAS-btz RNAi/+; repo-GAL4/+*. Scale bar, 30 µm. (K, L) quantifications of the number of boutons normalized to muscle surface area (x1000) in the indicated genotypes. ****, p<0.0001; ***, p<0.001; **, p<0.01; ns, not significant by unpaired t-test. n=23 (*w*; *da-GAL4, btz/Df*), n=6 (*btz/+*), n=10 (*Df/+, UAS-BtzGFP/+*; *btz/Df*), n=8 (*btz/Df*), n=22 (*da>btz-GFP*; *btz/Df*), n=13 (*BG380-GAL4*; *btz/Df*; *BG380+24B>Btz-GFP*; *btz/Df*; *da-GAL4/+*), n=5 (*BG380>Btz-GFP*; *btz/Df*), n=11 (*24B-GAL4*; *btz/Df*; *da>btz RNAi*), n=12 (*24B>Btz-GFP*; *btz/Df*; *btz RNAi/+*), n=9 (*BG380+24B-GAL4*; *btz/Df*), n=15 (*BG380-GAL4/+*), n=14 (*BG380>btz RNAi*; *repo>btz RNAi*), n=16 (*mhc-GAL4/+*), n=28 (*mhc>btz RNAi*), or n=21 (*repo-GAL4/+*). NMJ size is reduced in *btz* mutants and rescued by expressing *UAS-btz* with *da-GAL4* or with the combination of the *BG380-GAL4* and *24B-GAL4* drivers, but not with any single driver. Expressing *UAS-btz RNAi* with *da-GAL4* or in motor neurons with *BG380-GAL4*, muscles with *mhc-GAL4* or glia with *repo-GAL4* also reduces NMJ size.

To determine in which cell types Btz functions to promote NMJ growth, we used RNAi to knock it down in motor neurons, muscles or glia. Depleting Btz from motor neurons with *BG380-GAL4* (an insertion in the *futsch* gene) (Sanyal, 2009), from muscles with *myosin heavy chain* (*mhc*)-*GAL4* or from glia with *reversed polarity* (*repo*)*-GAL4* all caused a similar reduction in synaptic bouton number normalized to muscle surface area (Fig. 3H-J, L), indicating that Btz is required in all three cell types. Consistent with this conclusion, we were unable to rescue NMJ growth in *btz* mutants by providing wild-type Btz only to motor neurons, or to both muscles and glia using *24B-GAL4* (an insertion in the *held out wings* (*how*) gene) (Zaffran et al., 1997) (Fig. 3E, F, K). However, expressing *UAS-Btz-GFP* in all three cell types by combining *24B-GAL4* with *BG380-GAL4* rescued NMJ size to the same extent as ubiquitous Btz expression (Fig. 3G, K).

### Btz controls NMJ size independently of the EJC

To test whether Btz regulates NMJ size through the EJC, we compared the effects of knocking down other EJC subunits in motor neurons to the effect of knocking down *btz*. We found that in contrast to the reduction in bouton number caused by expressing *btz* RNAi with *BG380-GAL4* (Fig. 4B, I), expressing *mago* RNAi or *eIF4AIII* RNAi with the same driver significantly increased the number of boutons relative to muscle surface area when compared to controls (Fig. 4C, D, I). We also used our *Btz-WT* and *Btz-HD* transgenes to determine whether this function of Btz requires the residues that interact with the EJC. We found that the reduced NMJ size of *btz* mutants was rescued equally well by both transgenes (Fig. 4E-I). These data indicate that unlike other known functions of Btz, its role in promoting normal synapse growth is not mediated by the EJC.

**Figure 4:**
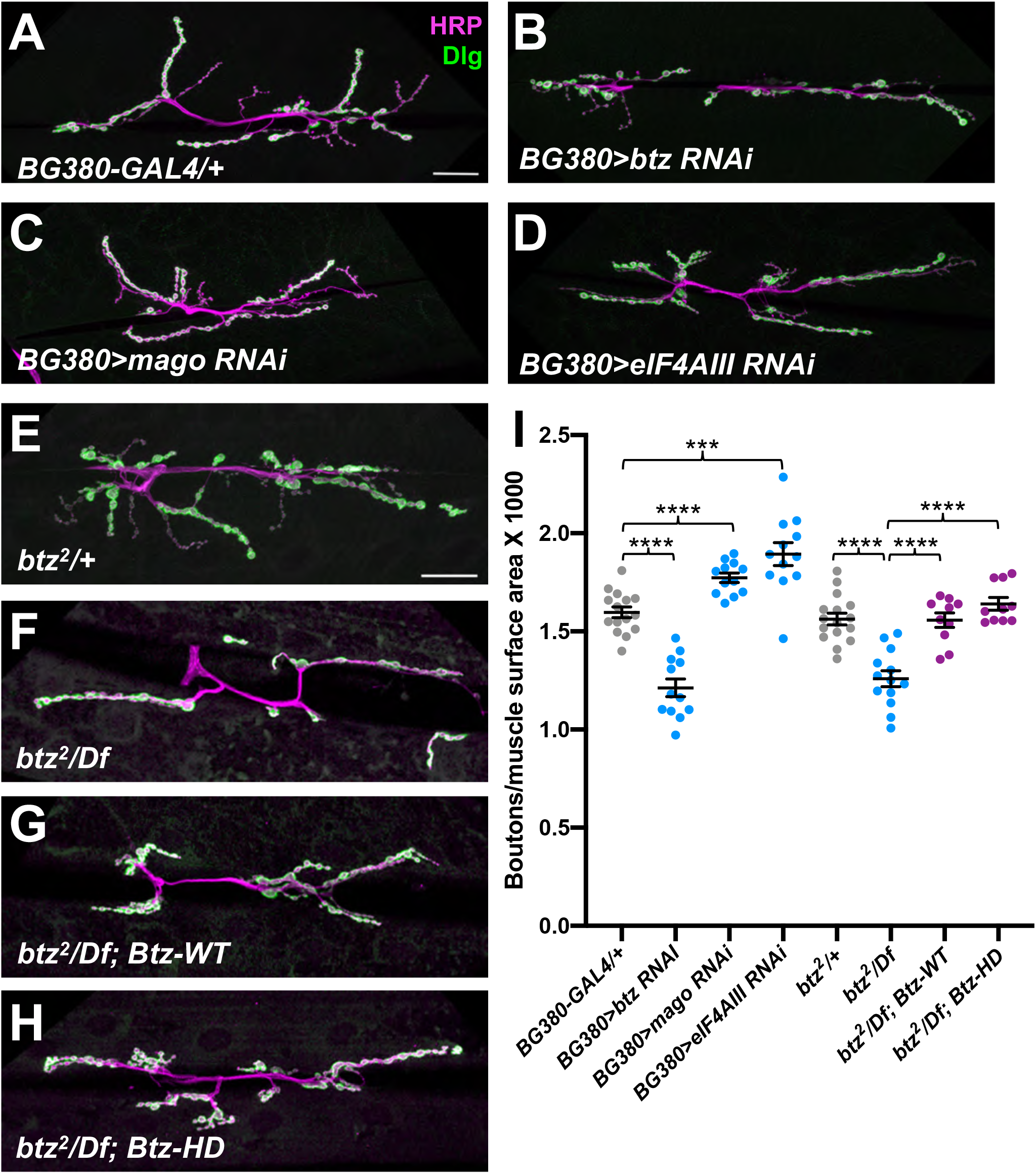
Btz controls NMJ size independently of the EJC. (A-H) confocal images of the NMJ on larval muscles 6 and 7 in segment A3, stained with anti-HRP (magenta) and anti-Dlg (green). (A) *BG380-GAL4/+*; (B) *BG380-GAL4/+*; *UAS-btz RNAi/+*; (C) *BG380-GAL4/+*; *UAS-mago RNAi/+*; (D) *BG380-GAL4/+*; *UAS-eIF4AIII RNAi/+*; (E) *btz*^*2*^*/+*; (F) *btz*^*2*^*/Df(3R)BSC497*; (G) *Btz-WT*; *btz*^*2*^*/Df(3R)BSC497*; (H) *Btz-HD*; *btz*^*2*^*/Df(3R)BSC497*. Scale bars, 30 µm (shown in A for A-D and in E for E-H). (I) quantifications of the number of boutons normalized to muscle surface area (x1000) in the indicated genotypes. ****, p<0.0001; ***, p<0.001 by unpaired t-test. n=15 (*BG380-GAL4*), n=12 (*BG380>btz RNAi*; *BG380>mago RNAi*; *BG380>eIF4AIII RNAi*), n=16 (*btz/+*), n=13 (*btz/Df*), or n=10 (*Btz-WT*; *btz/Df*; *Btz-HD*; *btz/Df*). NMJ size is reduced by knocking down *btz* in motor neurons, but slightly increased by knocking down *mago* or *eIF4AIII*. The size deficit in *btz* mutants is equally well rescued by the wild type or EJC interaction-defective *btz* transgenes.

### Btz acts through Dawdle signaling to control NMJ size

We next wished to identify targets of regulation by Btz that could explain its effect on NMJ size. We therefore examined gene expression in dissected central nervous systems or carcasses (consisting predominantly of body wall muscles) of *btz*^*2*^*/Df(3R)BSC497* third instar larvae compared to heterozygous *btz*^*2*^*/+* controls. From an RNA-Seq analysis of three biological replicates, we found 220 genes that showed significantly altered expression (q<0.05) in *btz* mutant larval carcasses and 103 that showed significantly altered expression in the central nervous system (Table S1). Some of these expression changes are likely to be due to the presence of other tissues in some samples, as they are genes that are highly expressed in tissues such as the salivary gland, fat body, or testis. In addition, genes located within *Df(3R)BSC497* showed an approximately 50% reduction in expression due to the absence of one wild-type copy. Removing these genes from the list left 80 potential Btz targets in muscle and 33 in the CNS (Table S1). One of the genes that was significantly downregulated in both the CNS and carcass samples in *btz* mutants was *daw*, which encodes an Activin homologue that has been implicated in the control of NMJ growth (Ellis et al., 2010; Fuentes-Medel et al., 2012). In addition, two genes that were upregulated in *btz* mutant carcasses, *larval translucida* (*ltl*) and *faulty attraction* (*frac*), encode proteins that can regulate the activity of TGF-β and BMP homologues in other contexts (Miller et al., 2011; Szuperak et al., 2011). We confirmed the changes in *daw, ltl* and *frac* expression by quantitative RT-PCR (Fig. 5A). Consistent with a reduction in signaling by members of the TGF-β family (Fuentes-Medel et al., 2012; Sulkowski et al., 2016), phosphorylation of the downstream factor Mothers against Dpp (Mad) was reduced at *btz* mutant NMJs (Fig. 5B, C, Fig. S1A, B). Mad phosphorylation was rescued by both the *Btz-WT* and *Btz-HD* transgenes (Fig. S1C, D), confirming that Btz regulates TGF-β signaling independently of the EJC. Loss of *btz* did not affect Mad phosphorylation in the ventral nerve cord (Fig. S1E, F), which is regulated by muscle-derived Gbb (Sulkowski et al., 2014; Sulkowski et al., 2016), indicating that its effect is limited to local TGF-β signaling at the synapse.

**Figure 5:**
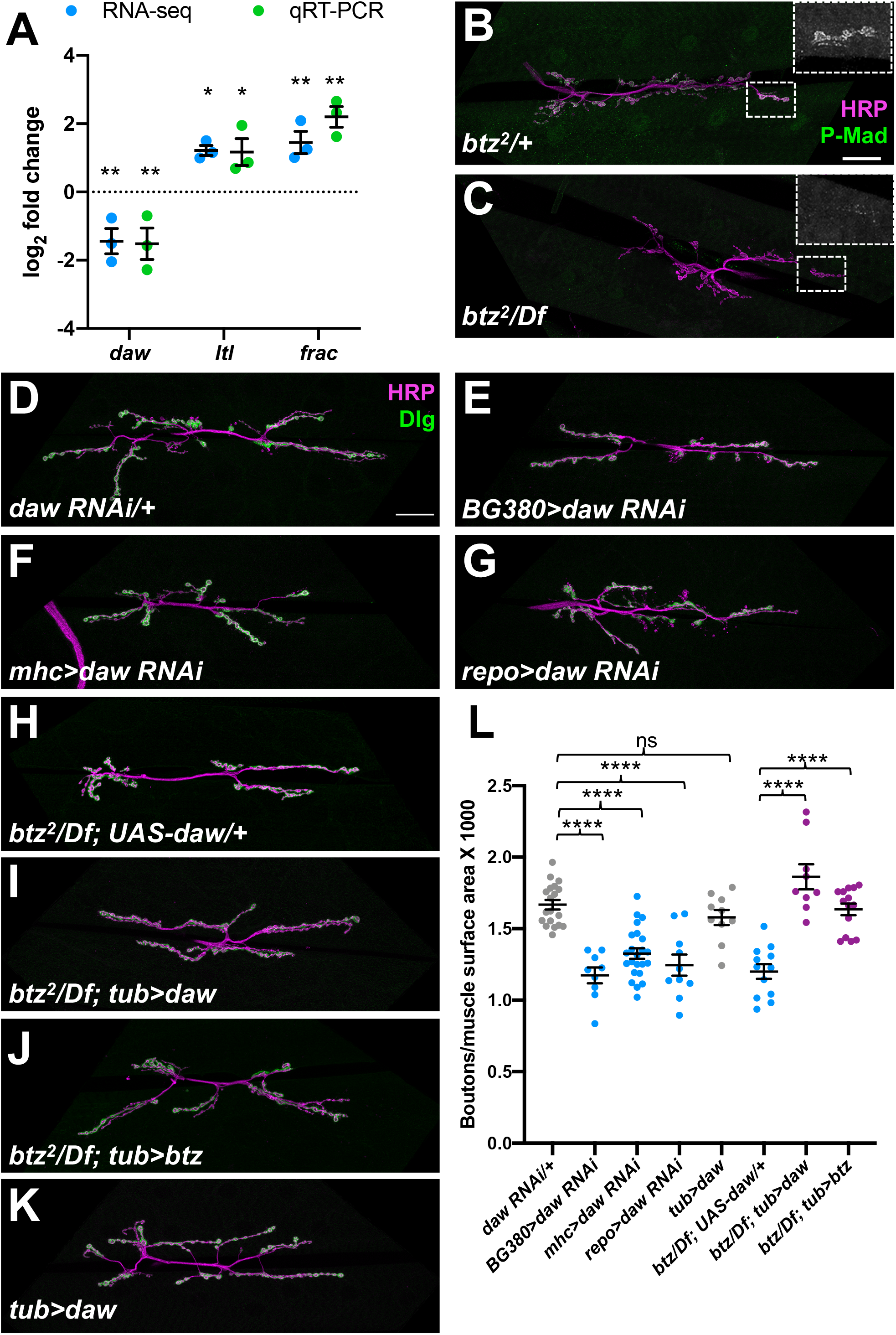
Daw signaling mediates the effects of Btz on NMJ size. (A) log_2_ fold changes in *daw, ltl* and *frac* mRNA in *btz*^*2*^*/Df(3R)BSC497* larval carcasses compared to *btz*^*2*^*/+*, measured by RNA-Seq (blue) or qRT-PCR (green). **, q<0.01; *, q<0.05. n=3 for each sample. (B-K) confocal images of the NMJ on larval muscles 6 and 7 in segment A3, stained with anti-HRP (magenta) and anti-P-Mad (green in B, C) or anti-Dlg (green in D-K). (B) *btz*^*2*^*/+*; (C) *btz*^*2*^*/Df(3R)BSC497*. Insets show enlargements of the boxed regions stained with anti-P-Mad. Synaptic P-Mad is lost in *btz* mutants. (D) *UAS-daw RNAi/+*; (E) *BG380-GAL4/+*; *UAS-daw RNAi/+*; (F) *mhc-GAL4/UAS-daw RNAi*; (G) *UAS-daw RNAi/+*; *repo-GAL4/+*; (H) *UAS-daw/+*; *btz*^*2*^*/Df(3R)BSC497*; (I) *tub-GAL4/UAS-daw*; *btz*^*2*^*/Df(3R)BSC497*; (J) *tub-GAL4/UAS-btz*; *btz*^*2*^*/Df(3R)BSC497*; (K) *tub-GAL4/UAS-daw*. Scale bars, 30 µm (shown in B for B-C and in D for D-K). (L) quantifications of the number of boutons normalized to muscle surface area (x1000) in the indicated genotypes. ****, p<0.0001; ns, not significant by unpaired t-test. n=19 (*UAS-daw RNAi/+*), n=9 (*BG380>daw RNAi*; *btz/Df; tub>daw*), n=23 (*mhc>daw RNAi*), n=10 (*repo>daw RNAi*; *tub>daw*), n=12 (*btz/Df*; *UAS-daw*), or n=14 (*btz/Df*; *tub>btz*). Knocking down *daw* in motor neurons, muscles or glia reduces NMJ size. Expressing *UAS-daw* with *tub-GAL4* rescues NMJ size in *btz* mutants as efficiently as expressing *UAS-btz*, but has no significant effect on NMJ size in a wild-type background.

*daw* was previously suggested to act in muscles and/or glia to regulate NMJ size (Ellis et al., 2010; Fuentes-Medel et al., 2012). We used RNAi to knock it down in muscles, motor neurons and glia, and found that like Btz, it was required in all three cell types for normal synaptic bouton numbers (Fig. 5D-G, L). To test whether Daw could mediate the effect of Btz on synapse growth, we expressed a *UAS-daw* transgene (Serpe and O’Connor, 2006) in *btz* mutant larvae. Ubiquitous expression of *daw* with *tubulin* (*tub*)-GAL4 rescued the *btz* mutant phenotype as effectively as ubiquitous expression of *btz* (Fig. 5H-J, L). Importantly, overexpressing *daw* with *tub*-GAL4 in wild-type larvae did not increase synaptic bouton number (Fig. 5K, L), indicating that it specifically restores a function that is lost in *btz* mutants. *daw* overexpression also partially rescued the lethality of *btz* mutants (Table 1). These results suggest that Btz regulates NMJ growth primarily by promoting Daw signaling.

## Discussion

### Btz acts both within and independently of the EJC

In this study, we used a rescue transgene in which the eIF4AIII-interacting residues of Btz are mutated to demonstrate distinct roles for Btz both as a component of the EJC, and independently of the EJC. We found that Btz is necessary for the normal distribution of mitochondria within muscles, and this function requires it to interact with the EJC. Little is known about the regulation of this mitochondrial distribution, although it can be altered by mutations that disrupt mitochondrial fission or increase fusion (Chao et al., 2016; Han et al., 2012; Wang et al., 2016a; Wang et al., 2016b). Several proteins that have been found to interact with Btz in a mass spectrometry screen are involved in regulating the expression of mitochondrial RNAs, although Btz itself is not thought to be localized to mitochondria (Baggio et al., 2014; Bratic et al., 2011; Guruharsha et al., 2011). Since the effect of Btz on mitochondria is EJC-dependent, it may result from a change in the translation or stability of a transcript that regulates mitochondrial localization. Interestingly, *oskar* mRNA, which is localized by the EJC in the ovary (Palacios et al., 2004; van Eeden et al., 2001), encodes a protein that traps mitochondria at the posterior pole of the embryo through interactions with the actin cytoskeleton (Hurd et al., 2016). However, we did not detect significant levels of *oskar* transcripts in the larval carcass in our RNA-Seq experiment.

In contrast, the function of Btz in promoting NMJ synapse growth is independent of the EJC, requiring neither the interacting residues of Btz nor the other subunits of the complex. The increase in NMJ size when *mago* or *eIF4AIII* is knocked down suggests that the EJC may even antagonize this function, perhaps by reducing the concentration of free Btz. Although it is clear that the other EJC subunits can perform important functions independently of Btz (Ashton-Beaucage et al., 2010; Blazquez et al., 2018; Boehm et al., 2018; Haremaki and Weinstein, 2012; Hayashi et al., 2014; Malone et al., 2014; Mao et al., 2017; Michelle et al., 2012; Roignant and Treisman, 2010), there are few reports to date of Btz functions independent of the EJC.

Additional studies will be required to determine the mechanism by which Btz regulates the levels of *daw* and other mRNAs that contribute to NMJ growth. Although the crystal structure of the core EJC shows that eIF4AIII forms most of the contacts with mRNA, with Btz contacting only a single base (Bono et al., 2006), Btz has been reported to be capable of binding mRNA in vitro (Degot et al., 2004). Indeed, one of the few reports of a potentially EJC-independent function for this subunit is that MLN51 regulates the translation of some mRNAs that lack introns (Chazal et al., 2013). Alternatively, it is possible that another RNA-binding protein links Btz to the target mRNAs responsible for its function in NMJ growth. The *Drosophila* genome encodes 29 members of the DEAD-box family in addition to eIF4AIII, and several are known to function in the nervous system. In a mass spectrometry screen, Btz was isolated in complexes with RNA-binding proteins that include Syncrip, which regulates Gbb levels in muscle (Halstead et al., 2014), as well as CG4612, Spenito and Quaking-related 58E-3 (Qkr58E-3) (Guruharsha et al., 2011). We have not determined whether Btz directly interacts with the *daw, ltl* and *frac* mRNAs or whether it controls their levels through an indirect mechanism; for instance, *daw* expression in the fat body is sensitive to dietary and metabolic changes (Chng et al., 2014).

### Regulation of Daw signaling controls NMJ size

Our data identify Daw signaling as an important mediator of the effect of Btz on NMJ growth. *daw* expression is reduced in both muscle and CNS in *btz* mutants, and increasing *daw* levels is sufficient to restore normal NMJ size in the absence of *btz*. Daw was previously reported to be produced by muscles and glia and to act on muscles to promote NMJ growth (Ellis et al., 2010; Fuentes-Medel et al., 2012; Parker et al., 2006; Serpe and O’Connor, 2006). However, we find that both *btz* and *daw* are required in three distinct cell types, muscles, motor neurons and glia, for normal NMJ growth. As Daw is a secreted protein, the reason for this is not immediately clear. It is possible that the requirement is quantitative, and production of sufficient Daw requires its secretion from all three cell types. As overexpressing *daw* in a wild-type background does not increase NMJ size, growth does not appear to be proportional to Daw expression, but a threshold quantity of Daw may be required to trigger a fixed expansion of the NMJ. This would be consistent with the transient requirement for BMP signaling early in development for NMJs to reach their normal size (Berke et al., 2013). Alternatively, Daw might be differently processed or post-translationally modified in each cell type, or secreted as a heterodimer with different TGF-β family members or in a complex with different proteins. This type of mechanism has been reported for Gbb: presynaptically produced Gbb can be distinguished from the postsynaptically produced pool due to its trafficking by and release with the neuronal protein Crimpy (James et al., 2014).

In addition to reduction of Daw, increased expression of Ltl and Frac may contribute to the *btz* mutant phenotype. These proteins have been shown to regulate the availability of extracellular TGF-β family proteins in developing wing discs and motor neurons (Miller et al., 2011; Szuperak et al., 2011). Either a reduction or an increase of Frac or Ltl expression can negatively regulate TGF-β signaling (Miller et al., 2011; Szuperak et al., 2011), indicating that the correct dosage of these proteins is critical. They could affect the activity of Daw or of Gbb or other ligands such as Activin-β, which is released from motor neurons to regulate muscle size (Moss-Taylor et al., 2019), or Maverick and Myoglianin, which are produced by glia and regulate NMJ growth (Augustin et al., 2017; Fuentes-Medel et al., 2012; Sulkowski et al., 2014).

Another function of Activin signaling is to regulate glutamate receptor expression and distribution at the synapse (Kim and O’Connor, 2014), which in turn can affect synaptic levels of pMad (Sulkowski et al., 2014). This mechanism could explain why we observed a reduced level of synaptic pMad in *btz* mutants. Activin signaling similarly promotes synaptogenesis and synaptic function in the mammalian hippocampus (Hasegawa et al., 2014; Shoji-Kasai et al., 2007), and can boost axonal regeneration (Omura et al., 2015). Activin mRNA levels are regulated by neuronal activity (Andreasson and Worley, 1995; Dow et al., 2005), but the mechanism of this effect is not known. Our results raise the possibility that activity exerts its effects through a Btz-dependent but EJC-independent mechanism that impinges on Activin levels. The discovery that Btz can act separately from the EJC should prompt a broader exploration of its possible functions in neuronal granules (Fritzsche et al., 2013).

## Materials and Methods

### Fly stocks and genetics

To generate the *Btz-WT* and *Btz-HD* rescue transgenes, genomic DNA spanning from 27700111 to 27703445 on 3R was amplified by PCR and cloned into pattB-QF-hsp70. Mutations changing the H215 and D216 residues to A were introduced by PCR and Gibson cloning. The primers used to make *Btz-WT* were Btz5UTRF (tagcggatccgggaattgggGTCCAGCCGTCGATACCGTTTTC), Btz5UTRR (cacccttttcTTCCTGCCCAGTGGCCG), eGFPlinkerF (tgggcaggaaGAAAAGGGTGGGCGCGC), eGFPlinkerR (acgcctgcacTCACGTGGACCGGTGCTTG), Btz3UTRF (gtccacgtgaGTGCAGGCGTACGCCGG), and Btz3UTRR (tagaggtaccctcgagccgcAACTTCGTGTTAAACGTTTATTTAAATTCAATGATCGATTAGTTTTTGCAAATACATTTCTCTC). To make *Btz-HD, Btz-WT* was used as a template with the primers Btz5UTRF, BtzHDAAR (cgtgctgcAGACCAGCGATCACCGCC), BtzHDAAF (gctggtctGCAGCACGATTCGAGGCC) and Btz3UTRR. The transgenes were injected by BestGene and integrated at the VK2 *attP* site. *UAS-btzGFP* was generated by amplifying the coding region of *btz* cDNA LD31454 by PCR and inserting it into pTWG (*Drosophila* Genomics Resource Center) by Gateway cloning. The plasmid was injected into embryos and integrated at random locations on the second and third chromosomes. Ovary phenotypes were examined in germline clones generated by crossing *Btz-WT* or *Btz-HD*; *FRT82, btz*^*2*^/*SM6-TM6B* females to *hsFLP*/*Y*; *FRT82, ovo*^*D*^/*TM3* males and heat-shocking the offspring for 1 h at 37°C during the pupal stage. Other stocks used were *w*^*1118*^ (BL#3605), *Df(3R)BSC497* (BL#25001), *G14-GAL4* (Aberle et al., 2002), *C57-GAL4* (Budnik et al., 1996), *da-GAL4* (BL#55850), *BG380-GAL4* (BL#42736), *24B-GAL4* (BL#1767), *mhc-GAL4* (BL#55133), *repo-GAL4* (BL#7415), *tub-GAL4* (Lee and Luo, 2001), *UAS-btz RNAi* (BL#30482), *UAS-mago RNAi* (VDRC#28132), *UAS-eIF4AIII RNAi* (VDRC#108580), *UAS-blw RNAi* (VDRC#34664), *UAS-daw RNAi* (BL#34974) *and UAS-daw* (Parker et al., 2006).

### Immunohistochemistry

To examine larval NMJs, 50 first instar larvae were collected on a grape juice agar plate and incubated at 25°C until they reached the third instar larval stage. Larval fillets were prepared by pinning the larvae to silicone plates, dissecting them in ice-cold Ca^2+-^free HL3 saline (pH=7.4), fixing in 4% formaldehyde in PBS for 15 min at room temperature, and permeabilizing with 0.2% Triton X-100 in PBS. Adult ovaries were dissected in PBS, fixed in 4% formaldehyde in PBS for 30 min on ice, and permeabilized with 0.2% Triton X-100 in PBS. Both tissue types were stained with primary antibody overnight at 4°C. The following antibodies were used: rabbit anti-Vasa (1:5000; (Gilboa and Lehmann, 2004)), chicken anti-GFP (1:400; Life Technologies A10262), mouse anti-Discs large (Dlg) (1:50; 4F3 from the Developmental Studies Hybridoma Bank (DSHB)), mouse anti-ATP5A (1:500, Abcam 15H4C4), and rabbit anti-P-Mad (Persson et al., 1998; Sulkowski et al., 2014). Tissues stained with primary antibodies were washed three times for five minutes each with 0.2% Triton X-100 in PBS and incubated with fluorescently labeled secondary antibodies for two hours at room temperature. The primary antibodies were visualized with corresponding secondary antibodies conjugated to Alexa Fluor-488 or Alexa Fluor-633 (1:200, Jackson ImmunoResearch). The neuronal membrane was visualized with Alexa Fluor-488, Alexa Fluor-633, or TRITC-conjugated anti-horseradish peroxidase (HRP) (1:200; Jackson ImmunoResearch). The muscle cells were visualized either using the background signal of the other antibodies used to stain the NMJ or with TRITC-conjugated phalloidin (1:5000; Abcam). The stained tissues were mounted in Fluoromount-G (SouthernBiotech). Samples were imaged with a Leica SP5 or SP8 confocal microscope using a 63x oil objective. Images were captured with a resolution of 1024 x 1024 pixels and processed in ImageJ and Adobe Photoshop. The images shown are z projections of confocal stacks acquired from serial laser scanning or xz projections of these stacks (Fig. 2).

### Cell culture and Western blotting

S2 cells were transfected with Btz-WT or Btz-HD using Effectene (Qiagen) according to the manufacturer’s protocol. After 3 days, cells were boiled in Laemmli buffer and run on a 15% SDS-PAGE gel. Gels were transferred to nitrocellulose membranes (Biorad) and were blocked overnight with TBST (20 mM Tris pH 7.5/150 mM NaCl/0.2% Tween-20) supplemented with 10% low-fat milk. Membranes were incubated with TBST + 10% milk supplemented with antibodies for 1 hour at room temperature. Blots were washed with TBST for an hour and incubated with HRP-conjugated secondary antibodies (1:2000, Jackson Laboratories) for another hour. Blots were developed with enhanced chemiluminescence (Pierce). Antibodies used were chicken anti-GFP (1:10,000, Aves GFP-1010) and mouse anti-β-tubulin (1:5000, Sigma T4026).

### Electron microscopy

EM samples were processed as described (Ramachandran and Budnik, 2010) with some modifications. Briefly, body-wall muscles of third-instar larvae were dissected in Jan’s saline containing 128mM NaCl, 2mM KCl, 4mM MgCl_2_, and 35.5mM sucrose in 5mM HEPES buffer (pH 7.2) with 0.1M CaCl_2_. The body-wall muscles were pinned on a silicone dish using insect pins and fixed with modified Trump’s fixative containing 4% paraformaldehyde, 1% glutaraldehyde, and 2mM MgCl_2_ in 0.1 M sodium cacodylate buffer (pH 7.2) at room temperature for 30 minutes and then overnight at 4°C. Fixed muscles were rinsed with 0.1 M sodium cacodylate buffer and post-fixed with 1% OsO4 in 0.1 M cacodylate buffer, followed by block staining with 1% uranyl acetate aqueous solution overnight at 4°C. The samples were rinsed in water, dehydrated in a graded ethanol series, infiltrated with propylene oxide/Spurr mixtures and finally embedded in Spurr resin (Electron Microscopy Sciences, PA USA). 70nm ultra-thin sections were cut and mounted on 200 mesh copper grids and stained with uranyl acetate and lead citrate. Imaging was performed on an electron microscope (CM12, FEI, Eindhoven, The Netherlands) at 120 kV, and recorded digitally using a camera system (Gatan 4k x 2.7K) with Digital Micrograph software (Gatan Inc., Pleasanton, CA).

### Quantification

The surface of muscles 6 and 7 was outlined and the enclosed area was quantified in ImageJ. The numbers of synaptic boutons were counted manually. To quantify mitochondrial localization, ImageJ was used to measure fluorescent intensity in random line scans taken from xz-projections of ATP5A-stained muscle. The highest peak, at the dorsal surface of the muscle, was normalized to the second-highest peak within the muscle.

All quantifications were carried out blind. Statistical significance between each genotype and the controls was determined by two tailed Student’s *t-*test, with Welch’s correction when the variances were significantly different. Each figure legend has details about sample sizes, precision measures, statistical analysis, and definitions of significance thresholds. No outliers were excluded.

### RNA extraction

Third instar larvae were dissected in 4°C DEPC-treated 0.1M phosphate buffer (pH 7.4). Dissected central nervous systems and larval fillets were mechanically homogenized with a plastic pestle in 200 µl TRIzol (Invitrogen). Total RNA was extracted from the samples using TRIzol /chloroform extraction: the larval fillets were incubated with a total volume of 450 µl TRIzol for 5 minutes and centrifuged at 12,000 rpm for 10 minutes at 4°C. The resultant supernatant was incubated with 107 µl chloroform, shaken vigorously by hand for 15 seconds, incubated at room temperature for 10 minutes, and centrifuged at 10,000 rpm for 10 minutes at 4°C. Approximately 250 µl of the upper aqueous phase was transferred to a new tube and incubated with 267 µl isopropanol at room temperature for 10 minutes. The RNA was pelleted by centrifugation at 12,000 rpm for 10 minutes at 4°C. The extracted RNA was washed 2x with 70% ethanol and purified using RNeasy Purification Kits (Qiagen). The RNA was eluted in 100 µl RNAse-free water and further purified and concentrated by sodium acetate precipitation: 10 µl 3M sodium acetate and 440 μl 100% ethanol were added to the RNA solution and it was incubated at −80°C overnight. The RNA was pelleted by centrifugation at 12,000 rpm for 30 minutes at 4°C, and the pellet was washed 2x with 70% ethanol and resuspended in water.

### Quantitative reverse transcription polymerase chain reaction

The purified RNA was treated with RQ1 RNase-Free DNase (Promega). Reverse transcription was performed from 1 μg of total RNA using SuperScript™ II Reverse Transcriptase (Thermo Fisher Scientific). Quantitative reverse transcription polymerase chain reaction (qPCR) was carried out using 10 ng of cDNA and 100 nM of each primer pair with a Roche LightCycler 480 machine and LightCycler 480 SYBR Green I Master 2X (Roche, 04887352001). The PCR program was: 10 min at 95°C, 45 cycles of 95°C for 15 s and 60°C for 1 min. The primers used were AGACCATAGCCATCCAGTCG and GCAGGTGATTTGGATGAGGT for *daw*, AGCTACGGATGCAGTGGAAC and GCCGATGCTTAGGTAGGTGA for *ltl*, and CACCACCGATGGAAAAACTT and TCAAAACTGATCTTGGGTTCG for *frac*.

### RNA-Seq

Total RNA was isolated from larval carcasses or central nervous systems from each genotype in triplicate using Trizol (Invitrogen). RNA quality and quantity was assessed using the Bioanalyzer 2100 (Agilent Inc.). Library preparation and sequencing was carried out by the NYU Genome Technology Center. RNA-Seq library preps were constructed using the Illumina TruSeq RNA sample Prep Kit v2 (Cat #RS-122-2002), using 500 ng of total RNA as input, amplified by 12 cycles of PCR, and run on an Illumina 2500 (v4 chemistry), as paired-end reads of 50 bp. For each RNA-seq sample, sequence quality was assessed with FastQC (http://www.bioinformatics.babraham.ac.uk/projects/fastqc) and sequencing adapters were removed with Trimmomatic (Bolger et al., 2014). Cleaned reads were aligned to the *Drosophila* reference genome (dm3) with Tophat2 v2.1.1 1. The Picard CollectRnaSeqMetrics program (https://broadinstitute.github.io/picard/picard-metric-definitions.html#RnaSeqMetrics) was used to generate QC metrics including ribosomal RNA content, median per-gene coverage, bases aligned to intergenic regions, 5’/3’ biases, and the distribution of the bases within exons, UTRs, and introns. Per sample gene expression profiles were computed using Cufflinks v2.2.1 1 and the RefSeq genome annotation for the *Drosophila* reference genome dm3 (Celniker et al., 2002). For multi-sample comparison, Principal Component Analysis and hierarchical clustering were used to verify that the expression profiles of the sequenced samples clustered as expected by sample tissue and genotype. Differential gene expression was computed for various contrasts between genotypes with the Cufflinks protocol (Trapnell et al., 2012) with default thresholds. RNA-Seq data have been submitted to NCBI GEO (reference number GSE165971).

## Acknowledgements

We thank Kavita Arora, Thomas Hurd, Ruth Lehmann, Brian McCabe, Micaela Serpe, the Bloomington *Drosophila* Stock Center, the Vienna *Drosophila* Resource Center, the *Drosophila* Genomics Resource Center and the Developmental Studies Hybridoma Bank for fly stocks and reagents. We thank NYULH DART Microscopy Laboratory members Alice Liang, Chris Petzold and Kristen Dancel-Manning for consultation and assistance with transmission electron microscopy; this core is partially funded by NYU Cancer Center support grant NIH/NCI P30CA016087. Flybase provided invaluable information for this study. We are grateful to Hui Hua Liu, DanQing He and Ariel Hairston for technical assistance. The manuscript was improved by the critical comments of Maria Bustillo, Neha Ghosh, Ashley Jordan, and Hongsu Wang. This work was funded by NSF grant MCB-1051022 to J.E.T.

