## Supplementary Information for "An exon junction complex-independent function of Barentsz in neuromuscular synapse growth"

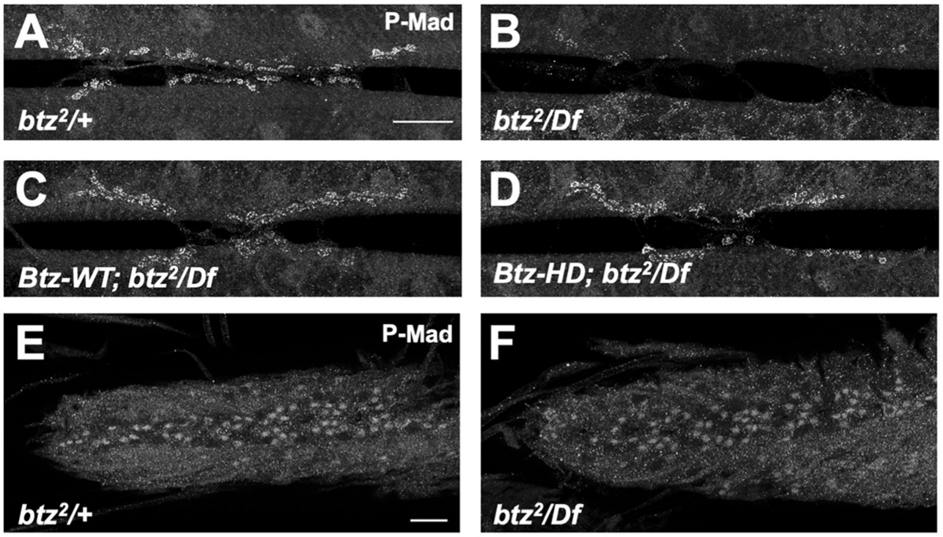


**Figure S1: Btz regulates P-Mad levels at the NMJ independently of the EJC.** Confocal images of the NMJ on larval muscles 6 and 7 in segment A3 (A-D) or the larval ventral nerve cord (E, F), stained with anti-P-Mad. (A, E) *btz*$^{2}$*/+*; (B, F) *btz*$^{2}$*/Df(3R)BSC497*; (C) *Btz-WT*; *btz*$^{2}$*/Df(3R)BSC497;* (D) *Btz-HD*; *btz*$^{2}$*/Df(3R)BSC497*. Scale bars, 20 μm. P-Mad is lost from the synapse but not from neuronal cell bodies in *btz* mutants, and is rescued by both the wild type and EJC interaction-defective *btz* transgenes.

**Table S1: Changes in gene expression in *btz*$^{2}$*/Df(3R)BSC497* compared to *btz*$^{2}$*/+* measured by RNA-Seq.** Sheet 1 shows changes in the larval CNS and sheet 2 in the carcass, which consists primarily of muscle. For each gene, the average reads per kb per million reads are given for the three samples in each of the two genotypes, as well as the $\log_2$ fold change and q value. A q value of <0.05 was considered significant. The genes highlighted in orange are those that are likely to show true *btz*-dependent expression changes. The others are either most highly expressed in a tissue other than muscle that may be present in some of the carcass samples, or fall within *Df(3R)BSC497* and show a two-fold reduction in expression in *Df(3R)BSC497* heterozygotes.
